# All-Optical Mapping Reveals Distributed Suppression of Cortical Sensory Responses After Optogenetic Silencing

**DOI:** 10.1101/2024.12.12.628207

**Authors:** Ambrosone Manuel, Montagni Elena, Resta Francesco, Ulivi Tommaso, Curti Lorenzo, Polverini Federica, Mazzamuto Giacomo, Mannaioni Guido, Masi Alessio, Pavone Francesco Saverio, Allegra Mascaro Anna Letizia

**Affiliations:** European Laboratory for Non-Linear Spectroscopy, Florence, Italy; Neuroscience Institute, National Research Council, Pisa, Italy; Physics and Astronomy Department, University of Florence, Sesto Fiorentino, Italy; National Institute of Optics, National Research Council, Sesto Fiorentino, Italy; Department of Neurosciences, Psychology, Drug Research and Child Health, University of Florence, Florence, Italy; Department of Health Sciences (DSS), University of Florence, Florence, Italy

**Keywords:** Optogenetics, all-optical, whiskers stimulation, inhibition, stGtACR2, barrel field cortex

## Abstract

**Significance:** We designed a novel all-optical tool to simultaneously silence neuronal activity at arbitrary sites on the dorsal cortex, and monitor the consequences of the manipulation. Optogenetic inhibition of primary sensory regions determined short and long-term dampening of the sensory response across a distributed cortical network.

**Introduction:** Many fundamental processes of brain computation, such as sensory perception and motor control, heavily rely on the mesoscopic dynamics of activity across the cerebral cortex. Manipulating mesoscale activity and observing its effects across multiple brain regions is crucial for understanding the causal link between cortical dynamics and behavior.

**Objective:** The goal of this study was to develop a novel all-optical system that allows inhibition of excitatory neurons while simultaneously monitoring cortical responses at arbitrary sites across the entire dorsal cortex of mice.

**Methods:** We combined wide-field imaging and optogenetics to create a mesoscale all-optical approach, enabling simultaneous monitoring and manipulation of cortical activity using light. Intravenous injection of two PHP.eB AAVs enabled the whole-brain co-expression of the red-shifted calcium indicator jRCaMP1b and the inhibitory actuator stGtACR2, with stable expression over several weeks. This system was calibrated, and the effects of inhibition on sensory responses were tested.

**Results:** Increasing laser power progressively reduced spontaneous activity at the site of irradiation. A single 5-second pulse on the barrel field cortex significantly decreased the amplitude of sensory-evoked responses, not only in the stimulated region but across the entire stimulated hemisphere.

**Conclusions:** This novel all-optical system enables targeted inhibition while concurrently monitoring mesoscale cortical activity. It provides insights into the dynamics of cortical circuits and offers a milestone for investigating the causal links between neuronal activity and behavior. Future research can use this tool to address sensory responsiveness impairments in neurological and neuropsychiatric disorders.

## 1. Introduction

Establishing a causal link between neural activity patterns across the entire brain and behavior is a major challenge in contemporary neuroscience. In the last decades, optical tools were increasingly adopted for disentangling this connection[1]. In the context of light-mediated control of neuronal activity, optogenetics demonstrated reversible depolarization or hyperpolarization of neurons through the illumination of light-gated ion channels with specific wavelengths[2]. Optogenetic tools, taking advantage of genetically encoded actuators for the activation or inhibition of neurons, hold significant promise for neuromodulation studies[3,4]. Recently, these tools have enabled precise neuromodulation by providing targeted inhibition of neuronal activity, allowing controlled perturbation of cortical responses to peripheral stimuli during active sensing at single cell resolution[5]. In this study, the authors leveraged the newly developed GtACR2, an anion-conducting channelrhodopsin from the cryptophyte *Guillardia theta* optimized for the in vivo silencing of neuronal activity with blue light (480 nm)[6–8]. Compared to other inhibitory actuators, GtACR2 shows positive photon-ion stoichiometry and good single-channel conductance[9]. Because of its improved light sensitivity, low axonal excitation, and high photocurrent, the soma-targeted variant (stGtACR2) is more effective for targeted silencing[9]. Due to its minimal responsivity above 560 nm wavelengths, it can be used in combination with red-shifted genetically encoded indicators, like RCaMPs, for simultaneous inhibition and imaging of cortical activity. RCaMP sensors are well-suited for tracking changes in calcium concentration, a direct parameter of neuronal activity[1,10]. They also provide several benefits when compared to the GCaMP family, including reduced photodamage, minimal hemodynamic interference, and increased maximal imaging depth[11,12]. Large-scale wide-field fluorescence imaging (WFFI) of red-shifted indicators is now commonly used for the measurement of cortex-wide neuronal activity[13–16]. By combining WFFI with optogenetic tools, it is now possible to measure and modulate large neuronal populations simultaneously using only light. Recently, this combination has been leveraged to perform large-scale all-optical monitoring and excitation of cortical activity[17–21]. Albeit stGtACR2 has been used in combination with red-shifted indicators to investigate small neuronal circuits[22], the consequences of local optogenetic inhibition on the cortical dynamics at the mesoscopic scale are largely unexplored. By leveraging WFFI in combination with the red-shifted calcium indicator jRCaMP1b and the blue-light actuator stGtACR2, we established a robust framework for investigating the large-scale effects of targeted cortical inhibition. Importantly, we validated the efficacy of our tool in reducing the cortical response to peripheral whiskers’ stimulation across the entire stimulated hemisphere.

## 2. Material and methods

A total of 13 female littermates C57BL/6 mice were used. All experiments were performed in accordance with the guidelines of the Italian Minister of Health (aut. n. 721/2020). All animals were housed in standard condition cages with a 12h light/dark cycle and food and water ad libitum. The experimental group included 8 mice while 5 mice were used as sham.

### 2.1. Virus injection

In order to induce the transfection of the excitatory neurons with the fluorescent calcium indicator jRCaMP1b and the optogenetic actuator stGtACR2, 8 mice were injected with a single retro-orbital injection of two different AAV-PHP.eB mixed. The final injected solution was prepared diluting in 112.5 µL of saline solution, 25 µL of indicator (ssAAV-PHP.eB/2-mCaMKIIα-NES_jRCaMP1b-WPRE-hGHp(A), 1.2x10E13 vg/ml) and 12.5 µL of actuator (ssAAV-PHP.eB/2-mCaMKIIα-stGtACR2_mCerulean-WPRE-bGHp(A), 1.7x10E13 vg/ml) obtained from the Viral Vector Facility, Zurich. Thus, each mouse received 3.0x10E11vg of indicator and 2.1x10E11vg of actuator. The sham group of animals (n =5 mice) were injected with a solution of 25 µL of indicator and 112.5 µL of saline.

### 2.2. Intact skull window surgery

This procedure was performed as described by Montagni and colleagues[23]. Briefly, 10 days post injection (dpi) mice were anesthetized with isoflurane (3% for induction, 1–2% for maintenance) and positioned in a stereotaxic apparatus (KOPF, model 1900). Ophthalmic gel (Lacrilube) was applied to protect their eyes, and body temperature was maintained using a warming pad. After administering local lidocaine, the skin and periosteum were removed and reference points were marked (bregma and lambda). A custom-made head bar was attached and secured with dental cement (Super Bond C&B – Sun Medical) posterior to lambda. A thin layer of transparent dental cement was applied to the exposed bone and let it dry for ∼10 minutes. Following surgery, mice were allowed to recover in a controlled environment with food and water available.

### 2.3. In vivo imaging setup and procedures

#### 2.3.1. Widefield fluorescence microscope

Imaging and optogenetic stimulation are performed with a custom-made wide-field setup based on our previous works[19,21]. Shortly, the jRCaMP1b excitation source was a red LED (595nm LED light, M595L3 Thorlabs, New Jersey, United State) with a bandpass filter (578/21 nm, Semrock, Rochester, New York, USA) and a dichroic mirror (606nm, Semrock, Rochester, New York, USA) directing light through an objective (TL2X-SAP 2X Super Apochromatic Microscope Objective, 0.1NA, 56.3 mm WD, Thorlabs) towards the skull. The stGtACR2 actuator was stimulated with a continuous wave laser (lambda = 473 nm, OBIS 473 nm LX 75mW, Coherent, Santa Clara, CA, USA) merged into the imaging path via a dichroic beam splitter (FF484-Fdi01-25 336, Semrock, Rochester, New York, NY, USA). The system used a random-access scanning head with two acousto-optic deflectors (DTSXY400, AA Opto-Electronic, Orsay France) and a laser beam with an approximate size of 200 μm. Fluorescence emitted by jRCaMP1b was captured through a bandpass filter (630/69, Semrock, Rochester, New York, USA) and imaged onto a high-speed CMOS camera (Orca Flash 4.0 Hamamatsu Photonics, NJ, USA) at 40 Hz with 512x512 pixels resolution that provided a 11.6x11.6 mm² field-of-view of the mouse cortex. To quantify stGtACR2 fluorescence across the entire dorsal cortex, we used a blue LED (415 nm, M415L4, Thorlabs, New Jersey, USA) coupled with a bandpass excitation filter (432/36 nm, Semrock, Rochester, New York, USA) and a dichroic mirror (425 nm, Semrock, Rochester, New York, USA). Emitted fluorescence was filtered using a bandpass emission filter (480/15 nm, Semrock, Rochester, New York, USA) and collected with the CMOS camera.

#### 2.3.2. Wide-field imaging in awake mice

Head-fixed imaging sessions were performed 20 and 30 dpi. Image stacks from each session were registered using a custom-made software. The custom software allows centering of the image while the mouse is under the microscope; this is obtained by overlaying a cross on the field of view (FOV) over the bregma landmark, which is marked by a black spot on the skull. In this way, each mouse in each recording session can be aligned based on the bregma position. Optogenetic calibration was performed on the first imaging session (20 dpi), while in the second imaging session (30 dpi), whiskers stimulation was performed. At each time point, the entire FOV was illuminated with a single pulse of the red and blue LEDs, in order to quantify the fluorescence of the two fluorophores over the entire dorsal cortex. LEDs light intensity was 4 mW/mm². For the functionality of the jRCaMP1b we also recorded a 3-minute imaging epoch of the resting state activity.

#### 2.3.3. Calibration of optogenetic irradiation

Laser stimulation patterns were generated using two orthogonally-mounted Acousto-optic Deflectors (AA Opto-electonic). A reference image of the FOV was used to target the laser beam on a selected cortex area (x=32.8mm on the right and y=10.3mm behind the bregma; barrel-field cortex: BFD). Single-pulse laser irradiation consisted of one pulse (10 ms ON) repeated for 10 times (15 s PRE-stim, 11 s POST-stim each session) at increasing laser powers (0.15 –0.32 – 0.82 –1.66 – 3.34 – 6.85 – 10.4 mW). 5-second laser irradiation consisted of 5 seconds of irradiation repeated for 10 times (15 s PRE-stim, 6 s POST-stim each session) at different laser power (0.15 – 0.32 – 0.82 mW). Optogenetic inhibition at different time durations was achieved by delivering laser irradiation of varying lengths (1-s, 2-s, 3-s, 4-s, and 5-s), repeated 10 times at a power of 0.82 mW (15 s PRE-stim, 6 s POST-stim each session).

#### 2.3.4. Unilateral multi-whisker stimulation

Single air puffs (60 ms long) were presented to the left whiskers in each imaging epoch (15 s PRE-stim, 11 s POST-stim). Whisker stimulation was performed using the Picospritzer III (General Valve™), with the air tube positioned approximately 2 cm away from the vibrissae. Whisker stimulation during irradiation consisted of 40 imaging epochs, where the air puff was delivered 3 seconds after the onset of laser irradiation. This design provided a sufficient baseline period for calculating the ΔF/F. To evaluate the long term effects of the irradiation a total of 40 imaging epochs of whisker stimulation were recorded before the irradiation phase (PRE), followed by 10 repetitions of 0.82 mW laser irradiation on the BFD. The irradiation pattern of each repetition consisted of 15 s of pre irradiation, 5-s irradiation, 6 s post. Additional 40 imaging epochs of whisker simulation were performed (POST), starting 1-2 minutes from the interruption of the irradiation.

### 2.4. Electrophysiology and confocal imaging

#### 2.4.1. In vivo multiunit spike recording

Forty days after injection (40dpi), electrophysiological signals were recorded in the SSp-bfd cortex (−1mm AP; -3mm ML) of 2 mice anesthetized with an ip injection of ketamine/xylazine 80/10 mg. For the entire duration of the experiments, the mice did not exhibit movements, and the breathing rate was ∼1 Hz. Parylene C-insulated tungsten microelectrode (A-M Systems) with an impedance of 2 MΩ were used. The electrode was advanced through a small hole in the skull into layer 5 (800 μm below the dura) using a motorized micromanipulator (EXFO Burleigh PCS6000). Signals were amplified with a 3000 AC/DC differential amplifier, sampled at 10 kHz, high-pass filtered at 0.1 Hz, and low-pass filtered at 10 kHz. A reference electrode was placed on the motor cortex. Traces were then band-pass filtered 200-1000 Hz to isolate fast potential transients reported as spikes of multiunit activity (MUA). For cross-talk evaluation, multiunit spike frequency was calculated in sequential 5-second time windows of LED ON/LED OFF (4 mW). For assessing the effect of 5-second laser illumination on neuronal activation, multiunit spike frequency was calculated in sequential 5-second time windows of laser ON/laser OFF (0.82 mW).

#### 2.4.2. Ex vivo electrophysiology

Brain slices (350 μm) containing the BFD were obtained using a vibroslicer (Leica VT1000S) in ice-cold carboxygenated cutting solution (Sucrose 206mM, Glucose 25mM, NaHCO3 25mM, KCl 2.5mM, NaH2PO4 1.25mM, MgSO4 3mM, CaCl2 1mM). After at least 1 hour in warm artificial cerebrospinal fluid (NaCl 130mM, KCl 3.5mM, NaH2PO4 1.25mM, NaHCO3 25mM, Glucose 10mM, CaCl2 1mM, MgSO4 2mM, 32-34 °C), slices were placed in a recording chamber under a Nikon Eclipse E600FN microscope. Electrophysiological recordings used borosilicate capillaries (Harvard Apparatus) filled with K+ gluconate solution (120mM K-Gluconate, 15mM KCl, 10mM HEPES, 1mM EGTA, 2mM MgCl2, 5mM Na2Phosphocreatine, 0.3mM Na2GTP, 4mM MgATP, pH 7.3), resulting in 3–4 MΩ resistance. Signals were sampled at 10 kHz and filtered at 2 kHz with an Axon Multiclamp 700B. The stGtACR2 silencing effect was assessed using whole-cell current clamp configuration, applying 4 seconds of a 150 pA depolarizing step, during which 1s of 470 nm LED stimulation was applied to activate the inhibitory channel. The inhibitory effect was quantified by measuring the firing rate (Hz) before, during, and after the LED stimulation.

#### 2.4.3. Ex vivo confocal imaging

Mice were perfused with 20–30 mL of 0.1 M PBS (pH 7.6) and 100 mL of 4% paraformaldehyde (PFA). Brain coronal slices (100 mm thick) were cut with a vibrating-blade microtome (Vibratome Series 1500—Tissue Sectioning System). Considering the BFD position, imaging was performed with a Nikon Eclipse TE300C2 laser confocal scanning microscope (Nikon) equipped with a Nikon Plan Apo 20x air objective. The setup was equipped with 408 nm and 561 nm lasers to simultaneously excite jRCaMP1b and stGtACR2 respectively. Emission filters were 472/10 nm and 630/69 nm.

### 2.5. Data analysis

All data analyses were performed in MATLAB (MathWorks), ImageJ (Fiji), and Origin.

#### 2.5.1. Quantification of in vivo cortical expression of jRCaMP1b and stGtACR2

To evaluate the in vivo expression, a ROI was selected on the jRCaMP1b image, covering the entire dorsal cortex, excluding the edges of the skull preparation. For both the indicator and the actuator, the expression was quantified as the mean fluorescence of the ROI at different time points (20 and 30 dpi).

#### 2.5.2. Optogenetic calibration

For each imaging epoch, ΔF/F was calculated by taking the baseline F as the mean of the signal before the optogenetic irradiation. Frames affected by skull autofluorescence induced by laser irradiation were removed. For single-pulse irradiation, 1 frame was removed; for 5-s laser irradiation, 200 frames were removed. Image resize to 256x256 was performed and 10x10 pixels ROI was selected taking account of the stimulation site. For single-pulse irradiation, peak amplitude and full width at half maximum (FWHM) were calculated. For 5-s laser irradiation, the average ΔF/F before and after irradiation was measured.

#### 2.5.3. Whiskers stimulation

For each imaging epoch, ΔF/F was calculated by taking the baseline F as the mean of the signal before the whiskers stimulation time point. Average activity maps were extrapolated from the mean of the three maximum-activity frames. Quantification of activated area was extrapolated from the binarized masks based on the threshold of Mean + 2SD. A total of 22 ROIs were then selected (11 ROI for each hemisphere, 10x10 pixels) based on the bregma coordinates and the stimulation site. For each ROI, we averaged the 40 epochs of sensory-evoked activity and we extracted a 4-s window (2s PRE-stim, 2s POST-stim) to evaluate the differences of single area activity before and after optogenetic irradiation. Each subject underwent correlation mapping, which involved calculating the Pearson’s correlation coefficient between the average signals extracted from each ROI and those of the other ROIs (during a 1-second window before and after whisker stimulus (ws)), and then averaging the results within a session. Fisher’s r-to-z transform was used to transform the averaged single-subject correlation maps, which were then averaged over all animals in a group. Next, the previously acquired group-related averaged maps were re-transformed to correlation values (r-scores). To display difference matrices, group differences were computed as r(AFTER)-r(BEFORE) of the averaged correlation values. Statistical significance of pairwise ROI correlations was assessed using two-sample t-Test with correction for multiple comparisons via the Benjamini–Hochberg false discovery rate method (q = 0.05).

#### 2.5.4 Ex vivo confocal imaging

The fluorescent neurons in the cortex from 3 mice were analyzed by an experienced observer to quantify the percentage of cells showing co-expression of both the indicator and the actuator.

### 2.6. Statistical analysis

Paired T-Test was used for the quantification of fluorescence, comparison between PRE- and POST-irradiation activated area and peak amplitude. Friedman test was used to compare the multiunit firing rate (Hz) before, during, and after the optogenetic inhibition. Wilcoxon Signed Ranks Test was used to compare the multiunit firing rate (Hz) before and during LED illumination. One-way repeated measures ANOVA followed by Bonferroni correction was performed for the quantification of single-pulse peak amplitude and FWHM. Two-way repeated measures ANOVA followed by Bonferroni correction was performed for the quantification of the average ΔF/F PRE- and POST-irradiation. Group-level ROI-based FC differences between before- and after-ws groups were assessed by means of one-way repeated measure ANOVA followed by Tukey correction. Errors are reported as Standard Error of Means otherwise specified in figure legends, *p < 0.05, ** p < 0.01, *** p < 0.001, **** p<0.0001.

## 3. Results

### 3.1. Experimental plan

In our study, we aimed to calibrate a novel all-optical system for simultaneous monitoring and inhibition of excitatory neurons. For this purpose, mice were co-injected in the retro-orbital sinus with a solution of two different PHP.eB viruses to induce the expression of the jRCaMP1b indicator and the stGtACR2 actuator at P30. After 10 days, intact skull window surgery was performed allowing optical access to the dorsal cortex. In vivo characterization is conducted from 20 days to 40 days after the injection. Finally, ex vivo electrophysiology and confocal imaging were accomplished (Fig. 1A). Imaging sessions were also performed in a sham group of mice expressing only jRCaMP1b (Supplementary Fig. 3A).

**Figure 1.**
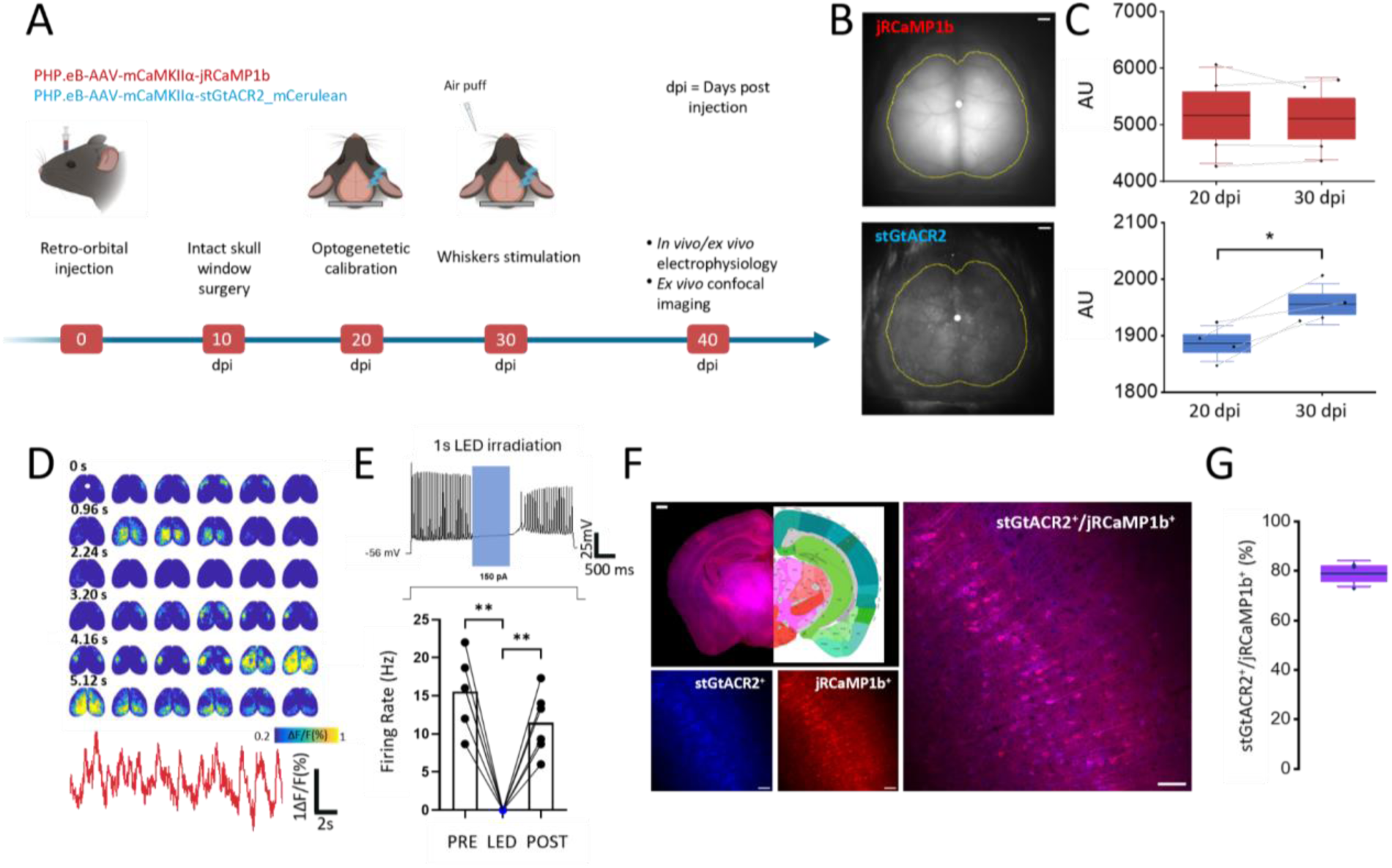
Experimental timeline and characterization of jRCaMP1b-stGtACR2 expression and functionality. **(A)** Experimental timeline with retro-orbital injection of both jRCaMP1b and stGtACR2, optical window surgery 10 days post injection (dpi), in vivo imaging sessions 20 and 30 dpi and in vivo electrophysiology 40 dpi. After in vivo experiments, ex vivo electrophysiology and confocal imaging analysis were performed. **(B)** Representative images of the field of view of jRCaMP1b (red) and stGtACR2 (blue). Yellow lines represent the selected ROI for analysis. White dot represents bregma, scale bar: 1mm. **(C)** in vivo quantification of absolute (dots) and average (line) jRCaMP1b (red) and stGtACR2 (blue) expression **(D)** Top: Representative image sequence of jRCaMP1b resting state cortical activity. White dot represents bregma, scale bar: 1mm. Bottom: Representative trace of jRCaMP1b activity averaged from the same ROI of panel B. **(E)** Top: Representative trace of neuronal activity before, during and after 1s optogenetic irradiation (in blue). Bottom: Bars graph of absolute (dots) and average (bars) measure of sEPSCs firing rate. **(F)** Upper left: Representative wide-field image of a single slice with related Allen Brain Atlas reference, scale bar: 1mm. Bottom left: Representative confocal images showing the neuronal expression of jRCaMP1b (red) and stGtACR2 (blue) in the barrel field cortex, scale bar: 100 µm. Right: Merge of the representative images, scale bar: 100 µm. **(G)** Quantification of the absolute (dots) and average (line) colocalization ratio stGtACR2^+^/ jRCaMP1b^+^.

### 3.2. Characterization of jRCaMP1b and stGtACR2 expression and functionality

To evaluate the efficacy and stability of the transfection, we quantified the in vivo expression of both jRCaMP1b and stGtACR2 over time. We obtained a high level of transfection of both viruses across the two imaging time points investigated (20, 30 dpi) (Fig. 1B-C) (red 20dpi: 5168±423 arbitrari unit (AU); red 30dpi: 5111±362 AU; blue 20dpi: 1886±16 AU; blue 30dpi: 1955±18 AU; n-mice=4; *p=0.02 Paired Sample t Test). Thanks to the homogeneous expression, we were able to measure changes in the calcium activity across the entire dorsal cortex (Fig. 1D). Additionally, we did not find any visible photobleaching in any 3-minute window of red light irradiation (Supplementary Fig.1A). For stGtACR2, ex vivo electrophysiological patch-clamp recordings showed single-cell inhibition during blue-light irradiation. Specifically, the firing rate during blue-light irradiation is significantly lower than the firing rates observed before and after the stimulation (Fig. 1E) (PRE: 10.37±1.286 Hz; LED: 0±0 Hz; POST: 7.63±1.133 Hz; n-mice=3, n-slices=6; PRE vs LED: **p=0.0011, LED vs POST: **p=0.0026, One-Way Anova, Tukey post hoc test). Using in vivo electrophysiology, we were able to verify the inhibition during the irradiation (Supplementary Fig.2 A) and the lack of stGtACR2 and jRCaMP1b cross-talk under single-photon excitation (Supplementary Fig.2 B). Histological analysis of brain slices revealed that both the indicator and actuator were largely expressed in the same neurons across the entire cortex (Fig. 1F). More than 75% of the cells expressing the jRCaMP1b indicator also expressed stGtACR2 in the BFD cortex (Fig. 1G) (78.9±3.03 %, n-mice=3), a co-expression pattern that was similarly observed in the motor and visual cortex (Supplementary Fig. 1B-C) (motor cortex : 82.3±0.54 %, visual cortex : 81.0±0.94 %, n-mice=2, n-slices=6). To sum up, we validate that a single retro-orbital injection of the mixed viruses induces stable expression and functionality of both the indicator and actuator.

### 3.3. Calibration of optogenetic inhibition of the barrel field cortex at different laser powers and durations

The effective expression and functionality of both actuator and indicator allowed us to simultaneously manipulate and monitor neuronal activity and test the consequences of irradiation at different laser powers and durations. We first calibrated 10-ms irradiation with increased laser powers (Fig. 2A). These brief pulses on the BFD activated a neuronal response immediately after the irradiation, which is proportional in amplitude to the laser power (Fig. 2B), in line with previous findings in vivo[9,24,25]. Specifically, low laser powers (0.15 mW and 0.32 mW) did not induce any measurable activation; however, we observed the emergence of post stimulus activity in the high range (from 0.82 mW to 10.4 mW), in which there is a significant increase in the peak amplitude of the stimulated area (Fig. 2C, Supplementary table 1) (0.15mW: 1.2±0.14%; 0.32mW: 1.6±0.13%; 0.82mW: 4.6±0.29%; 1.66mW: 7.3±0.44%; 3.34mW: 10.3±0.45%; 6.85mW: 12.3±0.68%; 10.4mW: 13.11±0.32%. n-mice=4, n-stim/mouse=10. See supplementary table 1 for significance). Significant differences in peak duration (FWHM) are observed when comparing high laser powers with the lower range. Within the higher range, the only significant difference in the FWHM of the post stimulus response was from 0.82 mW and 6.85 mW (Fig. 2D, Supplementary table 2) (0.15mW: 0.081±0.005s; 0.32mW: 0.0852±0.0007s; 0.82mW: 0.25±0.01s; 1.66mW: 0.198±0.017s; 3.34mW: 0.199±0.001s; 6.85mW: 0.201±0.002s; 10.4mW: 0.203±0.006s. n-mice=4, n-stim/mouse=10. See supplementary table 2 for significance). Finally, we calibrated the 5-second laser irradiation for three different laser powers (0.15 mW, 0.32 mW and 0.82 mW) (Fig. 2E). We demonstrated that irradiation effectively inhibits BFD activity during the 5-s post stimulus (Fig. 2F) (0.15mW PRE: 0.001±5.9*10^-4^ %; POST: -0.69±0.17%; 0.32mW PRE: 7.8*10^-4^ ±1.5*10^-4^ %; POST: -1.21±0,27%; 0.82mW PRE: 0.001 ±5.6*10^-4^ %; POST: - 2.17±0.41%; n-mice=4, n-stim/mouse=10). Within this interval, stronger inhibition was observed at the highest calibrated laser power (Fig. 2G). Importantly, this decrease of local activity is not observed in mice expressing only jRCaMP1b (Supplementary Fig. 3B-C) (0.15mW PRE: - 0.13±0.12 %; POST: 0.03±0.21%; 0.32mW PRE: -0.007±0.02 %; POST: 0.2±0.1%; 0.82mW PRE: -0.32±0.16%; POST: -0.2±0.3%; n-mice=3, n-stim/mouse=10). With the highest laser power (0.82 mW), we investigated the effect of inhibition at different durations (1 s, 2 s, 3 s, 4 s, and 5 s) (Fig. 2H-I). We found that even with the shortest duration (1 s), there is a significant reduction in local activity in mice injected with both jRCaMP1b and stGtACR2. Interestingly, changing the duration of the irradiation does not affect the extent of the inhibition (Fig. 2I) (1-s: PRE: 0.14 ± 0.07%; POST: −2.5 ± 0.8%; *p = 0.018; 2-s: PRE: 0.27 ± 0.09%; POST: −3.00 ± 0.57%; **p = 0.005; 3-s: PRE: 0.45 ± 0.06%; POST: −2.98 ± 0.39%; **p = 0.003; 4-s: PRE: 0.55 ± 0.03%; POST: −2.77 ± 0.44%; **p = 0.004; 5-s: PRE: 0.44 ± 0.07%; POST: −2.39 ± 0.34%; *p = 0.01; two-way repeated measures ANOVA, Bonferroni correction; n-mice=3, n-stim/mouse=10). In contrast, no significant changes were observed in mice only expressing jRCaMP1b (Supplementary Fig. 3D-E) (1-s: PRE: 0.02 ± 0.1%; POST: −0.08 ± 0.07%; 2-s: PRE: −0.03 ± 0.09%; POST: 0.17 ± 0.12%; 3-s: PRE: −0.06 ± 0.06%; POST: 0.05 ± 0.05%; 4-s: PRE: −0.04 ± 0.02%; POST: 0.05 ± 0.03%; 5-s: PRE: −0.006 ± 0.02%; POST: 0.12 ± 0.11%; n-mice=2, n-stim/mouse=10).

**Figure 2.**
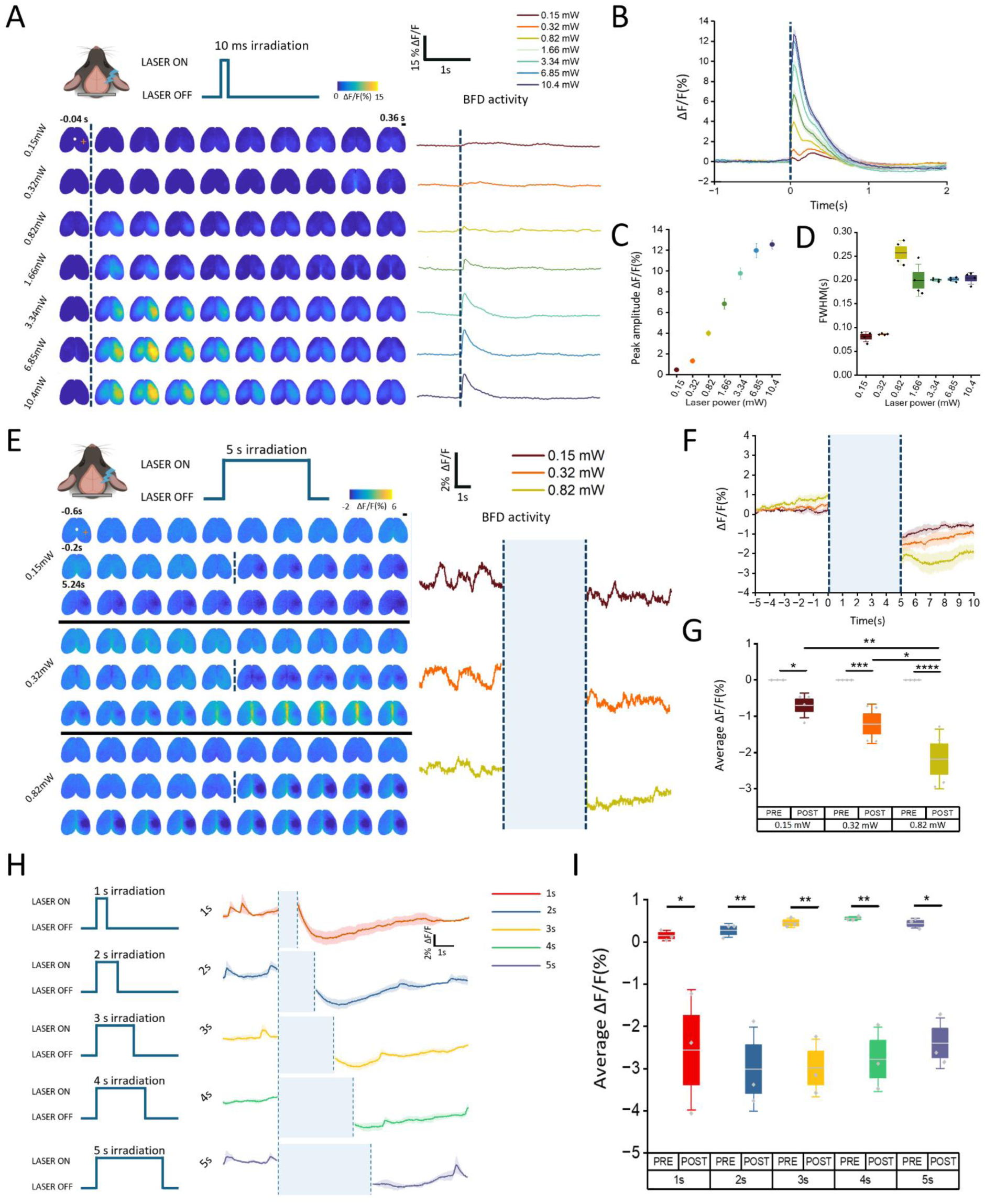
Optogenetic calibration of stGtACR2 stimulation at different laser power and durations. **(A)** Left: Representative image sequence of cortical activity before and after 10-ms blue-light irradiation on the barrel-field (BFD) at increasing laser power. Frame with optogenetic back-scattering is removed. Dashed line represents the stimulation time point. White dot represents bregma. Orange cross represents the stimulated site. Scalebar: 2mm. Right: Representative traces of BFD activity at increasing laser power. Dashed line represents the stimulation time point. **(B)** BFD activity before and after 10-ms blue-light irradiation (dashed line) at increasing laser power. **(C-D)** Quantification of peak amplitude (C) and FWHM (D) at increasing laser power **(E)** Left: Representative image sequence of cortical activity before and after 5-s blue-light irradiation on the BFD at increasing laser power. Frames with optogenetic back-scattering are removed. Dashed line represents the removed stimulation interval. White dot represents bregma. Orange cross represents the stimulated site. Scalebar: 2mm. Right: Representative traces of BFD activity at increasing laser power. Dashed lines represent the stimulation interval. **(F-G)** Averaged BFD activity before and after 5-s blue-light irradiation at different laser power (F) with relative quantification of average ΔF/F (G). Dashed lines represent the stimulation interval. **(H)** Averaged BFD activity before and after optogenetic stimulation at different durations. Dashed lines represent the stimulation interval. **(I)** Quantification of average ΔF/F 5s before (PRE) and after (POST) stimulation in the jRCaMP1b + stGtACR2 group.

### 3.4. Optogenetic inhibition silences the sensory-evoked response during laser irradiation

We tested if our optogenetic inhibition protocol could alter the excitability of the cortex. Therefore, we investigated if optogenetic inhibition of the BFD could modulate the cortical response to whiskers stimulation during irradiation. To this aim, three seconds after the onset of the irradiation at 0.82 mW, we delivered the whisker stimulation (Fig. 3A). In mice injected with both jRCaMP1b and stGtACR2, we did not observe a deflection of the signal compared to the baseline, whereas in mice expressing only jRCaMP1b, a clear peak of activation is observed following whisker stimulation (Fig. 3B,C). The difference in peak amplitude between these two groups is significant (Fig. 3D) (jRCaMP1b + stGtACR2: 0.19±0,04%; jRCaMP1b only: 0.71±0,04%; **p= 0.002, Two sample T-Test; nmice_ jRCaMP1b + stGtACR2=4, nmice_ jRCaMP1b_only=2, n-stim=40). These results suggest that optogenetic silencing was able to inhibit the insurgence of a sensory-evoked response during irradiation.

**Figure 3.**
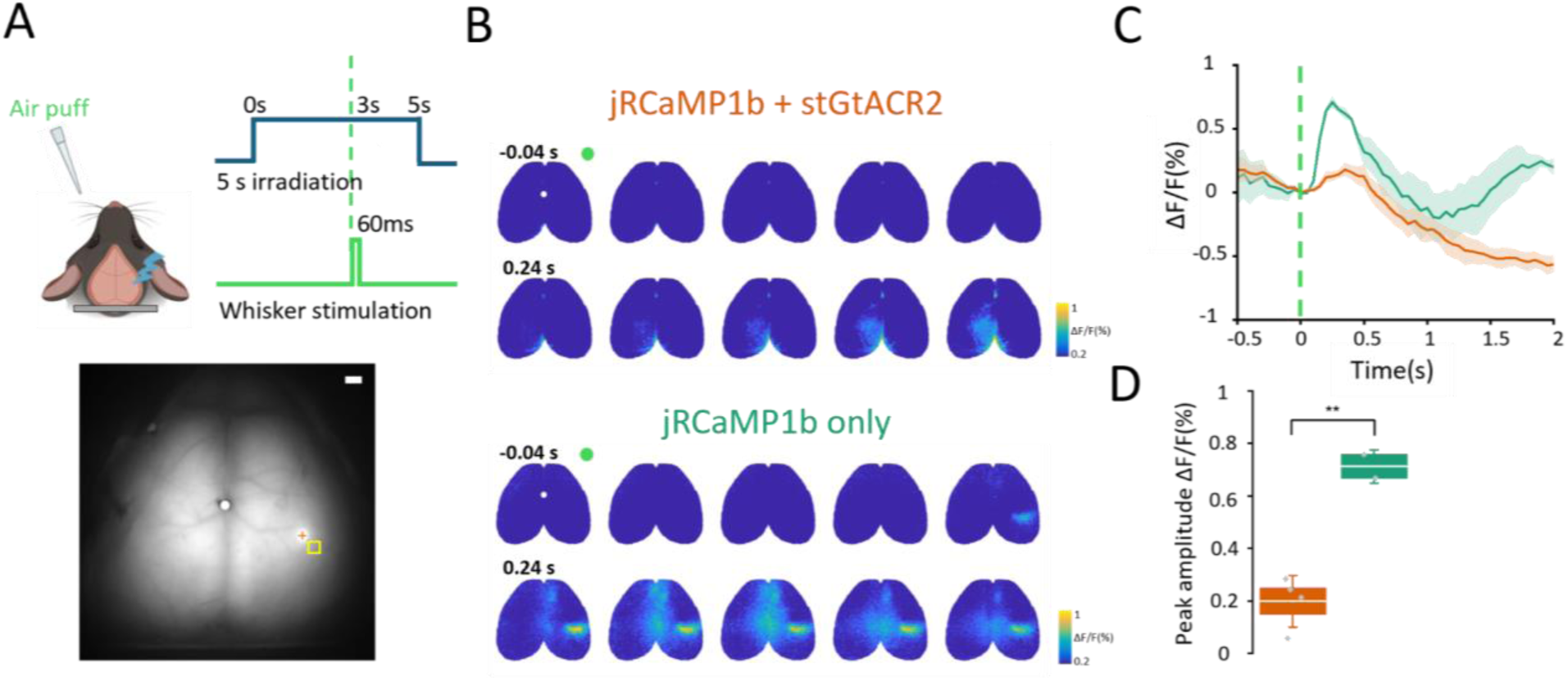
Whiskers stimulation during laser irradiation. **(A)** Top. Experimental paradigm with 5-s laser irradiation (blue) and whiskers stimulation (green). Bottom. Field of view indicating irradiation site (orange cross) and ROI for the quantification ΔF/F (yellow square). **(B)** Representative activation sequence of sensory-evoked response to whiskers stimulation during 5-s laser irradiation in the jRCaMP1b + stGtACR2 group (top, orange) and the sham group (bottom, green). Green dot represents the whisker stimulus time point. White dot represents bregma.**(C)** Sensory-evoked activity of the BFD cortex during the 5-s laser in the jRCaMP1b + stGtACR2 group (orange) and the sham group (green). Dashed lines indicate the whiskers’ stimulation time point. **(D)** Quantification of the absolute (dots) and average (line) peak amplitude in the jRCaMP1b + stGtACR2 group (orange) and the sham group (green).

### 3.5. Sensory evoked responses remain inhibited after optogenetic silencing

We then tested if local optogenetic silencing resulted in a reduction of network excitability even after irradiation. We applied the 5-s irradiation with the highest tested power (0.82 mW) on the BFD and evaluated the sensory-evoked response following the inhibition (Fig. 4A-B). We demonstrated that 5-s laser irradiation significantly decreased the size of the sensory-evoked activated area (Fig. 4C-D) (PRE: 95.3±16.0 mm^2^; POST: 44.6±5.1 mm^2^; n-mice=8; **p=0.004, Paired Sample t Test). Next, we compared the average sensory-evoked peak amplitude of the ipsi (right) hemisphere pre- and post-irradiation, observing a trend toward decreased sensory-evoked response (Supplementary Fig. 4A-B) (ipsi PRE: 0.41±0.05%; ipsi POST: 0.31±0.07%; nmice=8, n stim=40; p=0,08, Paired Sample t Test). We then quantified the effects of the irradiation on the sensory-evoked activity across different cortical areas (Fig. 4E). Inhibition extends beyond the stimulated area, significantly reducing sensory-evoked peak amplitude in both the BFD and the posterior parietal cortex (PPC) of the stimulated hemisphere (Fig. 4F) (Ssp-bfd_R PRE: 0.59±0.08%; Ssp-bfd_R POST: 0.41±0.07%; **p=0.003; PPC_R PRE: 0.64±0.07%; PPC_R POST: 0.44±0.08%; *p=0.028; VISp_R PRE: 0.33±0.06%; VISp_R POST: 0.20±0.06%; VISa_R PRE: 0.73±0.10%; VISa_R POST: 0.46±0.07%; p=0.053; n-mice=8, n-stim/mouse=40). We tested the same approach in mice expressing only jRCaMP1b with the 0.82 mW 5-s irradiation on the BFD. In these mice, neither the size of the sensory-evoked activation area nor the peak amplitude in the BFD and PPC showed any decrease (Supplementary Fig. 3F-J) (Activated area: PRE: 125.8±6.1 mm^2^; POST: 118.1±8.2 mm^2^; n-mice=5; Peak amplitude: BFD PRE: 0.74±0.35%; BFD POST: 0.67±0.33%; PPC PRE: 0.65±0.17%; PPC POST: 0.66±0.19%; n-mice=5, n-stim/mouse=40). Given that laser irradiation disrupts the sensory-evoked response across different regions of the stimulated hemisphere, we investigated how this inhibition influences the functional connectivity between cortical areas by comparing the correlation 1s before and after whiskers stimulation. We found a significantly reduced Pearson correlation between the stimulated BFD and the PPC (Fig. 4G-I) (PRE: 0.82±0.02r; POST: 0.86±0.01r; *p=0.006, Paired Sample t Test, n-mice=13, n-stim/mouse=40) while in the irradiation condition, the FC of the stimulated BFD did not show significant differences (Fig. 4J-L) (PRE: 0.83±0.01r; POST: 0.86±0.01r, n-mice=8, n-stim/mouse=40). These findings highlight the local and global effects of optogenetic inhibition, demonstrating its potential for targeted modulation of cortical excitability and responsiveness to peripheral stimuli.

**Figure 4.**
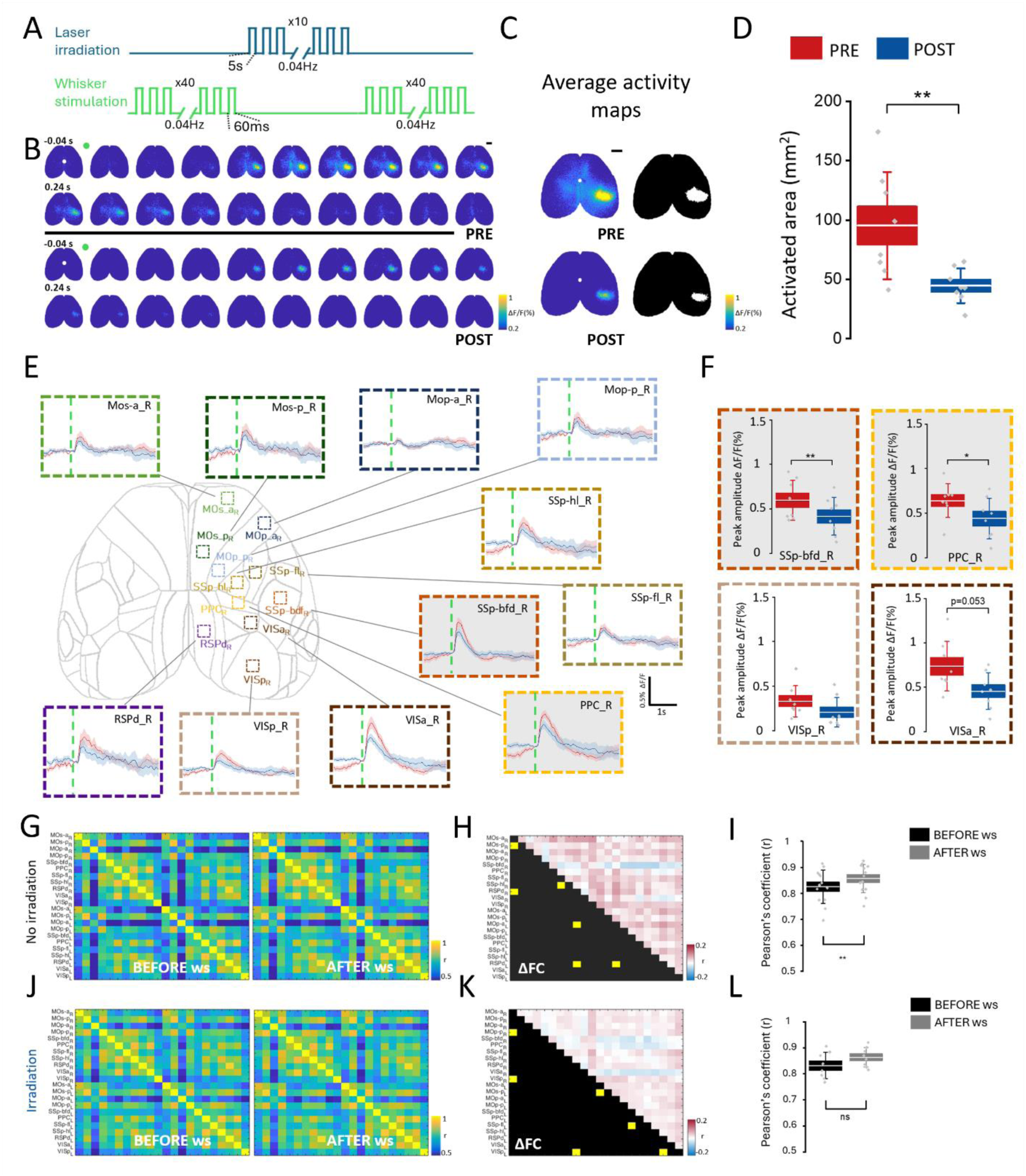
Effects of 5-second optogenetic stimulation on cortical sensory-evoked response. **(A)** Experimental setup with whisker stimulation(bottom) before and after 5-s laser irradiation (top). **(B)** Representative activation sequence of sensory-evoked response to whiskers stimulation before and after 5-s laser irradiation. Green dot represents the whisker stimulus time point. White dot represents bregma, scale bar: 2mm. **(C)** Left: Representative image of average activity maps before (PRE) and after (POST) 5-s laser irradiation with the correspondent binarized mask (right). White dot represents bregma, scale bar: 2mm. **(D)** Quantification of absolute (dots) and average (line) activated area before (red) and after (blue) optogenetic irradiation **(E-F)** Cortical parcellation with the relative sensory-evoked activity (E) before (red) and after (blue) optogenetic inhibition; quantification of representative areas peak amplitude (F) Dashed lines indicate the whiskers stimulation time point. Grey wallpaper indicates significant areas. **(G)** No-irradiation condition: Pairwise Pearson’s correlation coefficients visualized as average matrices of baseline- (left) and evoked-activity (right). **(H)** No-irradiation condition: Difference matrix of functional connectivity (FC) PRE- and POST-whisker stimulation (ws). Red and blue indicate POST hyper- or hypo-connectivity, with significant differences in yellow (*p<0.05; n-mice=13, 40 stimulations). **(I)** No-irradiation condition: BFD-PPC connectivity quantification PRE- (black) and POST-ws (grey). **(J)** Irradiation condition: Pairwise Pearson’s correlation coefficients visualized as averaged matrices of baseline- (left) and evoked-activity (right). **(K)** Irradiation condition: Difference matrix of FC PRE- and POSTt-ws. Red and blue indicate POST hyper- or hypo-connectivity, with significant differences in yellow (*p<0.05; n_mice=8, 40 stimulations). **(L)** BFD-PPC connectivity quantification PRE- (black) and POST-ws (grey).

## 4. Discussion

In the present study, we demonstrated the realization of a novel all-optical approach that allows simultaneous silencing and visualization of optogenetic inhibition effects at the mesoscale.

This was achieved by a single intravenous injection of two PHP.eB AAV driving whole-brain expression of both the optical indicator and actuator. This enables targeted silencing at arbitrary locations and monitoring of the resulting activity across an extended network spanning the entire dorsal cortex. To the best of our knowledge, this approach is novel. Previous all-optical studies have typically relied on intracranial injection of AAVs [26,21] or loading of a voltage sensitive dye in transgenic mice expressing ChR2 [17,27]. Other large-scale methods, such as electrophysiology or voltage imaging, are constrained by the limited number of recording sites— reducing spatial resolution—or by low signal sensitivity, respectively. In contrast, our method offers both high sensitivity and high spatial resolution, allowing for reliable, single-trial observations with fine spatiotemporal detail. Our results reveal that sensory-evoked responses are suppressed not only during optogenetic silencing but also for several minutes afterward. Crucially, the ability to detect this prolonged dampening in remote regions—such as the posterior parietal and visual cortices—was made possible only through our novel approach.

Our findings reveal how localized cortical inhibition influences sensory processing and interregional connectivity, addressing critical questions about the role of cortical dynamics in behavior.

We employed a single systemic injection of a mixed solution of two PHP.eB viruses to induce the expression of jRCaMP1b and stGtACR2 in mice. This approach ensured homogeneous and stable expression over several weeks (Fig. 1, Supplementary Fig. 1), avoiding the parenchymal damage associated with invasive microinjections reported in previous studies [28–30]. Unlike prior optogenetic tools, which often led to unintended excitatory effects due to off-target stimulation [24,25], the specific design of stGtACR2 minimizes axonal activation while maintaining high efficacy at the soma level [9]. This specificity is crucial for interpreting the causal relationships between neuronal silencing and cortical dynamics. However, the increasing expression of stGtACR2 over the weeks (Fig. 1C) raises concerns about potential actuator toxicity. Importantly, we verified the absence of cross-talk between stGtACR2 and jRCaMP1b under single-photon excitation (Supplementary Fig. 2 B), extending earlier findings with two-photon excitation using the same combination of actuator and indicator [26].

In line with other studies [31,32], 10-ms irradiation triggered a rebound of local activity, with light-dose dependent amplitude (Fig 2A-D). Either an increase in intracellular chloride or a decrease in intracellular proton concentrations could be responsible for this rebound activity. Indeed, fast variations in intracellular ion levels may cause either an efflux of chloride or an inflow of protons following light offset, which can result in rapid membrane depolarization [24,25]. In line with this hypothesis, recent studies focused on single cell inhibition through stGtACR2 reported ‘paradoxical’ neuron local excitation [32,33]. However, our data corroborate previous evidence [9,34] demonstrating local silencing of neuronal activity during LED irradiation both ex vivo and in vivo, respectively with patch clamp on brain slices (Fig. 1E) and LFP recordings (Supplementary Fig. 2 A). Moreover, our results reveal that optogenetic inhibition using stGtACR2 leads to robust and graded suppression of spontaneous activity in the BFD up to 5s after the irradiation (Fig 2E-I). While the multiunit firing rate recordings reveal a return to the baseline level right after the irradiation (Supplementary Fig 2A), the calcium imaging signals show a slower return to baseline (Figure 2E-F). This discrepancy likely reflects the different origin of the signal: while the fluorescence signal originates specifically from excitatory neurons, which are selectively silenced by the illumination, the field potential largely reflects the activity of inhibitory neurons, which are not affected by the optogenetic manipulation.

During irradiation, the BFD activation was markedly reduced in mice expressing both jRCaMP1b and stGtACR2 compared to those expressing only jRCaMP1b (Fig. 3). In the post irradiation period, the 5-second optogenetic inhibition caused a significant reduction in both the spatial extent and amplitude of sensory-evoked responses to whisker stimulation (Fig. 4B-G). This suppression extended beyond the local BFD region, involving interconnected cortical areas, including the PPC, which is well-known as a segregated integrative hub for the sensory-related processing [35,36]. Our work builds on this foundation by showing that mesoscale inhibition not only disrupts local sensory processing but also alters the network-wide coordination of cortical areas. The modulation of sensory responses by optogenetic inhibition reflects the fundamental role of distributed cortical processing in the elaboration of a sensory input [37]. Time-resolved optogenetic inhibition of the BFD response has been previously demonstrated by activation of GABAergic circuits [38,5]. Conversely, here we silenced CaMKII-positive neurons. Our approach offers a complementary tool to investigate sensory responses while providing a large-scale strategy to describe cortical network function.

These results also support the idea that connectivity patterns are crucial for explaining the paths through which inhibition flows into distant brain circuits. Indeed, local suppression subsequently alters the FC of several cortical regions. Specifically, without irradiation, whiskers stimulation induces a reduction in the FC of the stimulated BFD, indicating a weaker correlation with surrounding cortical areas and a more localized response (Fig. 4H). This result is in line with previous findings on sensory-evoked FC [39]. Here, the FC between the BFD and the PPC of the stimulated hemisphere is significantly higher in the 1-s window after the whiskers stimulation (Fig. 4I). However, after optogenetic inhibition reduced the amplitude of the stimulus-evoked activity in the BFD, the correlation difference between pre and post stimulus becomes non-significant (Fig. 4L). This result highlights the strong connection between these two areas in sensory integration and underscores its importance for sensorimotor functions, in line with previous findings [35,40]. Moreover, this disruption of FC has implications for understanding the neural basis of sensory impairments seen in disorders such as autism spectrum disorder and schizophrenia, where atypical connectivity patterns are thought to underlie perceptual deficits [41].

In conclusion, by demonstrating that the expression of both the indicator and the actuator can be achieved with a single injection, we proved the feasibility of applying this all-optical tool in animal disease models, while avoiding the complications associated with cross-breeding of multiple transgenic mouse lines.

In future studies, our tool could be leveraged to reduce the cortical sensory response in mouse models with hypersensitivity to peripheral stimuli. It would allow not only to correct behavioral anomalies associated with altered perception but also to study the immediate and long-term consequences on the cortical (dys)function. In addition, the system developed here could be adapted for brain-machine interface applications, where fine monitoring of cortical activity is essential for closed-loop control of behavior. The insights gained from our work could help optimize neural prosthetics for sensory restoration or motor rehabilitation, paving the way for more effective treatments for sensory and motor deficits.

This study establishes a novel all-optical framework for investigating the effects of targeted inhibition on mesoscale cortical dynamics. By demonstrating both local and global effects of inhibition, we provide new insights into the functional organization of cortical networks. Our findings contribute to a growing body of literature that emphasizes the importance of distributed sensory processing and highlights the potential of optogenetic tools for advancing both basic and translational neuroscience.

## Disclosures

Authors declare no competing interests.

## Code, Data, and Materials Availability

The data that support the findings of this study are available on request from the corresponding author.

## Acknowledgements

This work has been funded by Telethon Seed Grant Spring 2022 PHEM (GSA22E006), the Italian Ministry of Universities and Research PRIN 2022 (2022YCTLPL), project THE Tuscany Health Ecosystem (ECS_00000017), MUR_ PNRR, Banca di Italia and Fondo di Beneficenza di Intesa Sanpaolo RICONSIN project (B/2021/0202), Ricerca Scientifica e Tecnologica della Fondazione Cassa di Risparmio di Firenze (CRF2020). In addition, this project has been supported by the framework of Eurobioimaging (ESFRI research infrastructure) - Advanced Light Microscopy Italian Node. Figures 1A,E, 2A, 2E, 3A and Supplementary Figure 2A were created using BioRender.

## Contribution

ALAM conceived the study.

ALAM, MA and EM designed and interpreted experiments and computational analyses.

MA, EM, TU, FP, LC and FR performed experiments and generated data.

GMaz developed the software for the recording experiment.

MA, EM and FR conducted computational analyses.

MA, EM and ALAM drafted the original version of the manuscript.

LC, FR edited the manuscript.

ALAM, FSP, AM, GMan contributed funding and resources.

All authors participated in the review of the manuscript.

## Declaration of generative AI and AI-assisted technologies in the writing process

During the preparation of this work the authors used ChatGPT to improve the syntax and the readability of the manuscript. After using this tool, the authors reviewed and edited the content as needed and took full responsibility for the content of the published article.

---

**Supplementary Figure 1.**
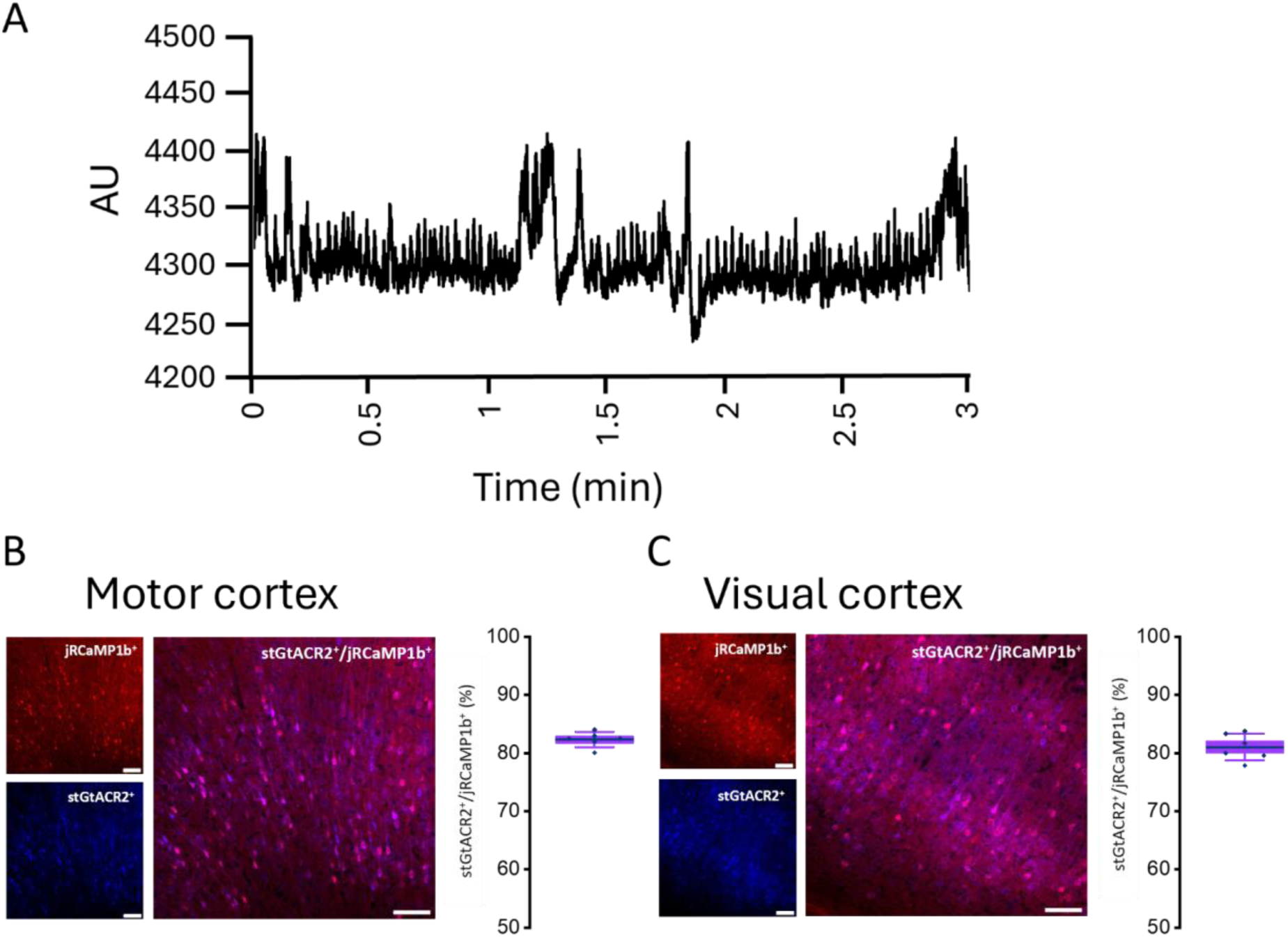
Raw fluorescence of jRCaMP1b and ex vivo expression in different cortical areas. **(A)** Representative raw trace of resting state fluorescence of the jRCaMP1b indicator within a 3 -minutes window. **(B)** Left: Representative confocal images showing the neuronal expression of jRCaMP1b (red) and stGtACR2 (blue) in the motor cortex with merge (purple), scale bar: 100 µm. Right: Quantification of the absolute (dots) and average (line) colocalization ratio stGtACR2^+^/ jRCaMP1b^+^. **(C)** Left: Representative confocal images showing the neuronal expression of jRCaMP1b (red) and stGtACR2 (blue) in the visual cortex with merge (purple), scale bar: 100 µm. Right: Quantification of the absolute (dots) and average (line) colocalization ratio stGtACR2^+^/ jRCaMP1b^+^.

**Supplementary Fig. 2.**
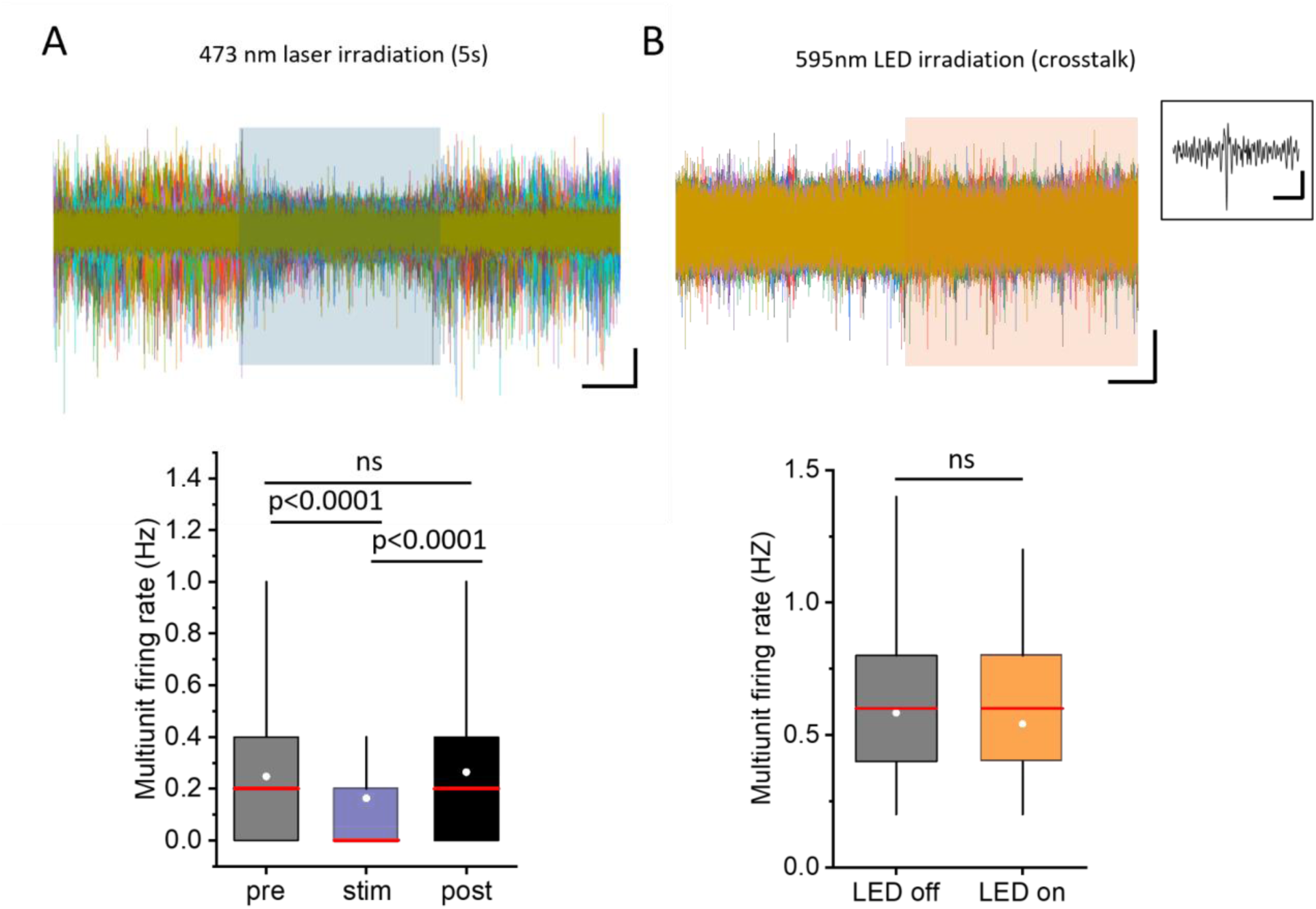
Electrophysiological validation of optogenetic inhibition and stimulation/imaging crosstalk. **(A)** Top: Representative traces showing multiunit spikes. The 5-second optogenetic stimulation period is represented by the light blue square. Scale bar: 100 µV / 1 s. Bottom: Quantification of the multiunit firing rate recorded during and post optogenetic stimulation (box = 25%-75%; whiskers = range within 1.5 IQR; red line = median; white circle = mean). Statistical test = Friedman test. **(B)** Top: Representative traces showing multiunit spikes. The LED-on period is represented by the light orange square. Scale bar = 50 µV / 1 s. Inset: Magnified view of a multiunit spike, scale bar 100 µV / 0.02 s. Bottom: Quantification of the multiunit firing rate recorded during imaging LED illumination (box = 25%-75%; whiskers = range within 1.5 IQR; line = median; circle = mean). Statistical test = Wilcoxon Signed Ranks Test.

**Supplementary Figure 3.**
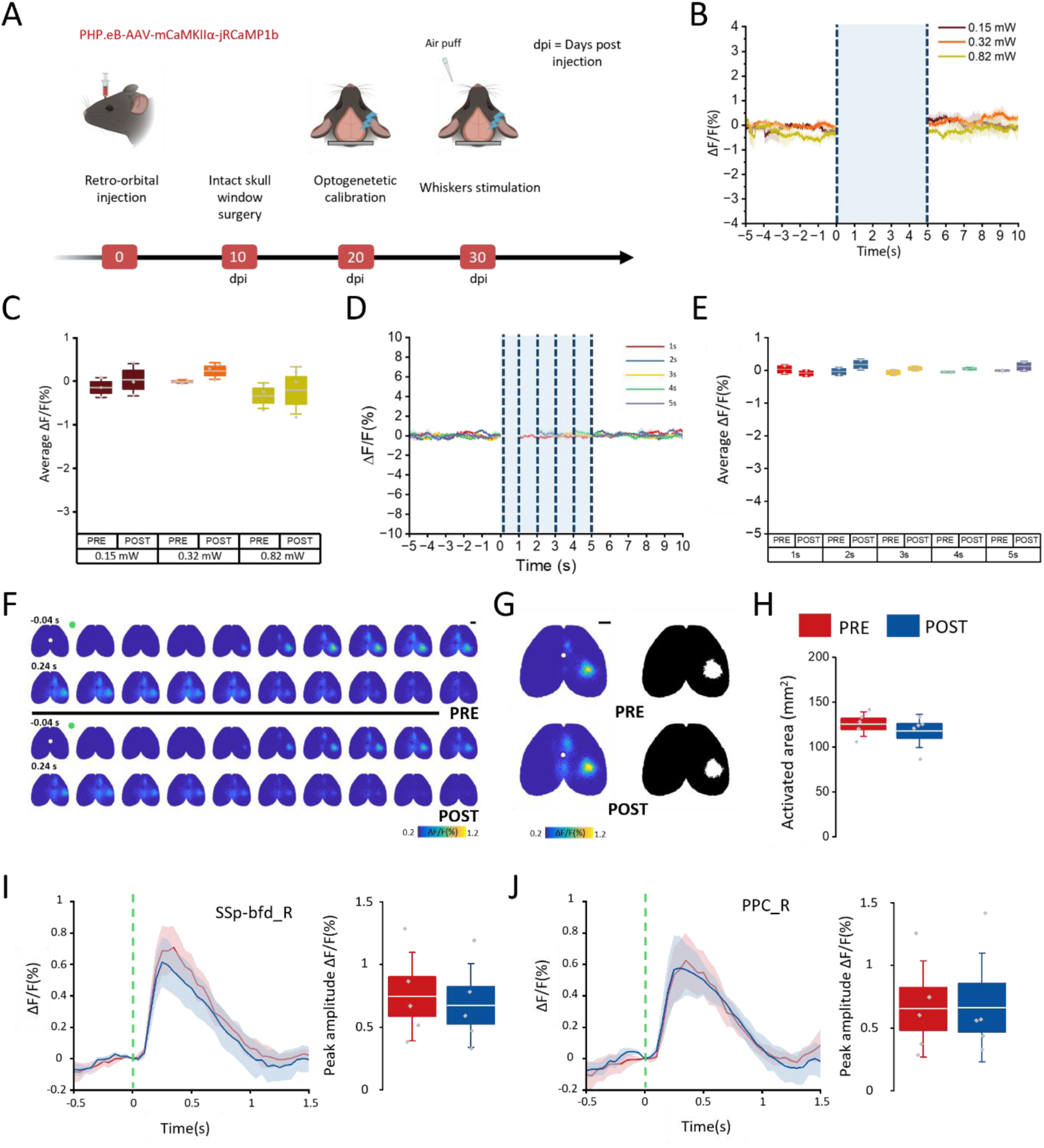
Effects of laser irradiation on mice not expressing stGtACR2. **(A)** Experimental timeline with retro-orbital injection of jRCaMP1b, optical window surgery 10 days after injection(dpi) and in vivo imaging session 20 dpi. **(B-C)** Averaged BFD activity before and after 5-s of optogenetic stimulation at different laser power (B) with relative quantification of average ΔF/F (C). Dashed lines represent the stimulation interval. **(D-E)** Averaged BFD activity before and after different durations of optogenetic stimulation (D) with relative quantification of average ΔF/F (E). Dashed lines represent the stimulation interval. **(F)** Representative activation sequence of sensory-evoked response to whiskers stimulation before and after 5-s laser irradiation. Green dot represents the whisker stimulus time point. White dot represents bregma, scale bar: 2mm. **(G)** Representative image of average activity maps before (left) and after (right) 5-s laser irradiation (top) with relative binarized masks (bottom). White dot represents bregma, scale bar: 2mm. **(H)** : Quantification of activated area before (red) and after (blue) optogenetic irradiation (PRE: 127.2±3.7 mm^2^; POST: 110.4±12.0 mm^2^; nmice=3). **(I)** Sensory-evoked activity of the BFD cortex before (red) and after (blue) optogenetic irradiation and quantification of peak amplitude. **(J)** Sensory-evoked activity of the PPC before (red) and after (blue) optogenetic irradiation and quantification of peak amplitude.

**Supplementary Figure 4.**
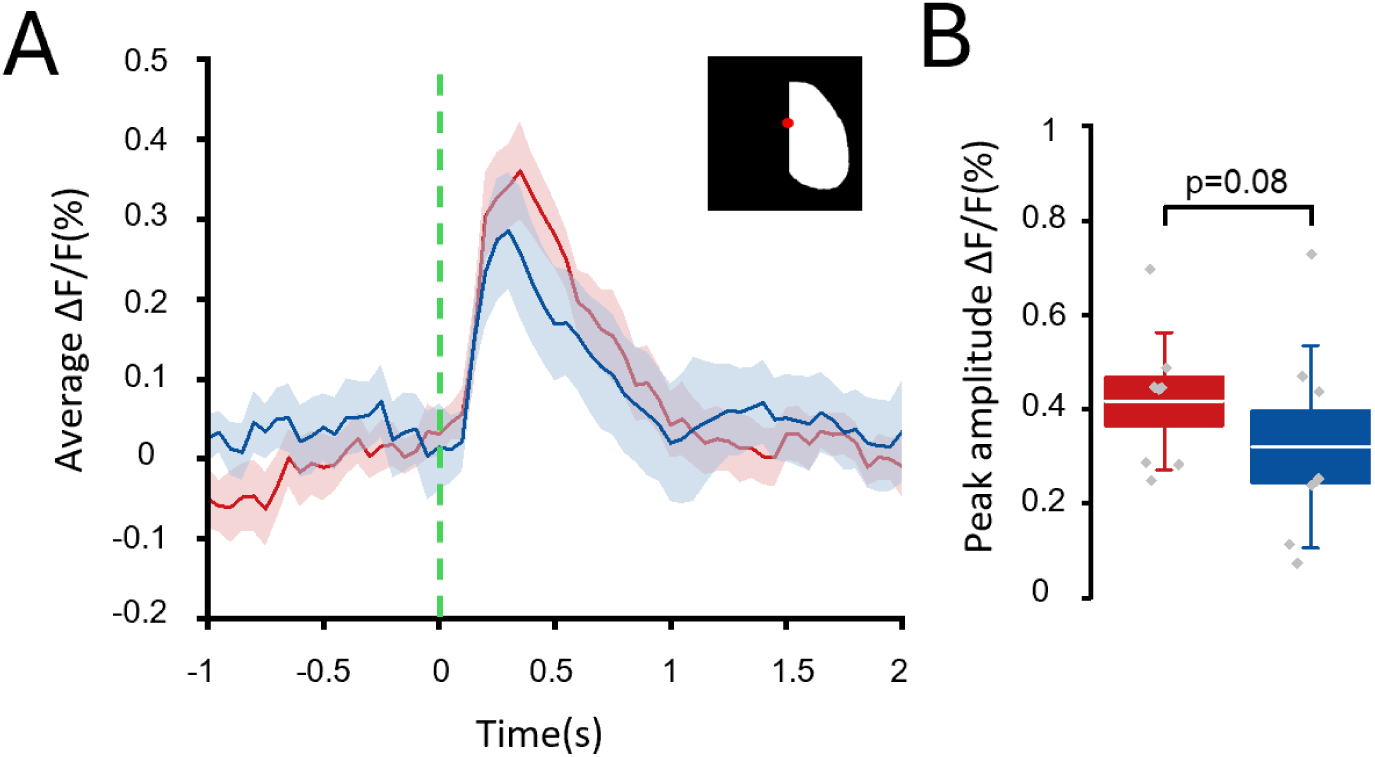
Effects of 5 second optogenetic stimulation on the stimulated hemisphere activity. **(A-B)** Sensory-evoked activity of the right (ipsi) hemisphere before (red) and after(blue) 5s optogenetic irradiation (A) with the correspondent quantification (B). Dashed line indicates the whiskers’ stimulation time point. Top panel refers to the ROI for the quantification of the ΔF/F, red dot indicates bregma.

**Supplementary Table 1.** One-way repeated measures ANOVA followed by Bonferroni correction for quantification of peak amplitude in single-pulse irradiation. Comparison between coupled powers with relative t Value, Prob>t and significance. *p < 0.05, ** p < 0.01, *** p < 0.001,****p<0.0001.

| **Powers** | **t Value** | **Prob>t** | **Sign.** |
| --- | --- | --- | --- |
| 0.15-0.32 mW | 1.05648 | 1 | ns |
| 0.15-0.82 mW | 9.44133 | 4.50896E-7 | **** |
| 0.15-1.66mW | 17.00494 | 3.26088E-11 | **** |
| 0.15-3.34mW | 25.63564 | 2.6817E-14 | **** |
| 0.15-6.85mW | 31.23952 | 8.23003E-16 | **** |
| 0.15-10.4mW | 33.2844 | 2.67772E-16 | **** |
| 0.32-0.82mW | 8.38485 | 2.61899E-6 | **** |
| 0.32-1.66mW | 15.94846 | 9.66218E-11 | **** |
| 0.32-3.34mW | 24.57916 | 5.60679E-14 | **** |
| 0.32-6.85mW | 30.18304 | 1.51202E-15 | **** |
| 0.32-10.4mW | 32.22792 | 4.74201E-16 | **** |
| 0.82-1.66mW | 7.5636 | 1.13334E-5 | **** |
| 0.82-3.34mW | 16.19431 | 7.46266E-11 | **** |
| 0.82-6.85mW | 21.79819 | 4.55306E-13 | **** |
| 0.82-10.4mW | 23.84307 | 9.54272E-14 | **** |
| 1.66-3.34mW | 8.6307 | 1.71799E-6 | **** |
| 1.66-6.85mW | 14.23458 | 6.48118E-10 | **** |
| 1.66-10.4mW | 16.27946 | 6.82943E-11 | **** |
| 3.34-6.85mW | 5.60388 | 5.38007E-4 | *** |
| 3.34-10.4mW | 7.64876 | 9.6961E-6 | **** |
| 6.85-10.4mW | 2.04488 | 1 | ns |

**Supplementary Table 2.** One-way repeated measures ANOVA followed by Bonferroni correction for quantification of FWHM in single-pulse irradiation. Comparison between coupled powers with relative t Value, Prob>t and significance. *p < 0.05, ** p < 0.01, *** p < 0.001, **** p<0.0001.

| **Powers** | **t Value** | **Prob>t** | **Sign.** |
| --- | --- | --- | --- |
| 0.15-0.32 mW | 2.44854 | 0.52118 | ns |
| 0.15-0.82 mW | 2.83438 | 0.23089 | ns |
| 0.15-1.66mW | 5.88381 | 3.0051E-4 | *** |
| 0.15-3.34mW | 6.01137 | 2.31225E-4 | *** |
| 0.15-6.85mW | 6.39831 | 1.05774E-4 | *** |
| 0.15-10.4mW | 6.24374 | 1.44221E-4 | *** |
| 0.32-0.82mW | 5.28292 | 0.00106 | ** |
| 0.32-1.66mW | 8.33236 | 2.86858E-6 | **** |
| 0.32-3.34mW | 8.45991 | 2.30081E-6 | **** |
| 0.32-6.85mW | 8.84685 | 1.19326E-6 | **** |
| 0.32-10.4mW | 8.69228 | 1.54764E-6 | **** |
| 0.82-1.66mW | 3.04943 | 0.14492 | ns |
| 0.82-3.34mW | 3.17699 | 0.10962 | ns |
| 0.82-6.85mW | 3.56393 | 0.04658 | * |
| 0.82-10.4mW | 3.40936 | 0.06565 | ns |
| 1.66-3.34mW | 0.12755 | 1 | ns |
| 1.66-6.85mW | 0.5145 | 1 | ns |
| 1.66-10.4mW | 0.35993 | 1 | ns |
| 3.34-6.85mW | 0.38694 | 1 | ns |
| 3.34-10.4mW | 0.23237 | 1 | ns |
| 6.85-10.4mW | 0.15457 | 1 | ns |

## Notes

### Competing Interest Statement

The authors have declared no competing interest.

### Summary of Updates

Figure 1, 2, 4 revised. Figure 3 added. Author added. Supplementary figures added.

